# Improving accuracy and precision of heritability estimation in twin studies through hierarchical modeling: Reassessing the measurement error assumption

**DOI:** 10.1101/2023.06.24.546389

**Authors:** Gang Chen, Dustin Moraczewski, Paul A. Taylor

## Abstract

**Introduction:** The conventional approach to estimating heritability in twin studies implicitly assumes either the absence of measurement error or that any measurement error is incorporated into the nonshared environment component. However, this assumption can be problematic when it does not hold or when measurement error cannot be reasonably classified as part of the nonshared environment.

**Methods:** In this study, we demonstrate the need for improvement in the conventional structural equation modeling (SEM) used for estimating heritability when applied to trait data with measurement errors. The critical issue revolves around an assumption concerning measurement errors in twin studies. In cases where traits are measured using samples, data is aggregated during preprocessing, with only a centrality measure (e.g., mean) being used for modeling. Additionally, measurement errors resulting from sampling are assumed to be part of the nonshared environment and are thus overlooked in heritability estimation. Consequently, the presence of intra-individual variability remains concealed. Moreover, recommended sample sizes are typically based on the assumption of no measurement errors.

**Results:** We argue that measurement errors in the form of intra-individual variability are an intrinsic limitation of finite sampling and should not be considered as part of the nonshared environment. Previous studies have shown that the intra-individual variability of psychometric effects is significantly larger than the inter-individual counterpart. Here, to demonstrate the appropriateness and advantages of our hierarchical linear modeling approach in heritability estimation, we utilize simulations as well as a real dataset from the ABCD (Adolescent Brain Cognitive Development) study. Moreover, we showcase the following analytical insights for data containing non-negligible measurement errors:

i. The conventional SEM may underestimate heritability.
ii. A hierarchical model provides a more accurate assessment of heritability.
iii. Large samples, exceeding 100 observations or thousands of twins, may be necessary to reduce imprecision.

**Discussion:** Our study highlights the impact of measurement error on heritability estimation and introduces a hierarchical model as a more accurate alternative. These findings have significant implications for understanding individual differences and improving the design and analysis of twin studies.

## 1 Introduction

As an indication of potential predictability, heritability is an important concept in assessing individual differences. As the proportion of trait variability ascribed to genetics, heritability offers a unique perspective for quantifying the role of genetics in complex traits (Downes and Turkheimer, 2022; Robette et al., 2022). Twins provide a hypothetically well-controlled scenario where genetics, environment, and their interaction can be statistically separated and apportioned.

### 1.1 Heritability estimation: ACE model and Falconer’s formula

Conventional twin studies are typically conceptualized with three hierarchies of data structure: individual, family, and zygosity. The individual measures are nested within families, which are further categorized as either monozygotic (MZ) or dizygotic (DZ) twins. A model can be formulated for a quantitative trait of interest that is measured at the individual level. In the popular ACE formulation (Maes, 2005; Downes and Matthews, 2020; Hunter, 2021), the trait data *y*_*i*(*f* (*z*))_ is expressed as the combination of latent components through the three indices of individual (*i* = 1, 2, …, *I*), family (*f* = 1, 2, …, *F*), and zygosity (*z* = MZ, DZ):

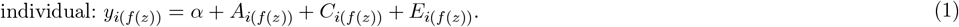

Each pair of parentheses indicates a nesting structure among the subscripts. The intercept *α* captures the overall trait effect at the population level. The acronym for the ACE model reflects the three latent sources of variability. *A*_*i*(*f* (*z*))_ represents the additive genetic effects, and *C*_*i*(*f* (*z*))_ represents the common or shared environmental effects. In addition, *E*_*i*(*f* (*z*))_ characterizes the unique or nonshared environmental effects.

The variances associated with the three latent components are crucial parameters in twin studies. One may make the following assumptions for two twins *i*_1_ and *i*_2_ within a family *f* of zygosity *z* (Arbet et al., 2020),

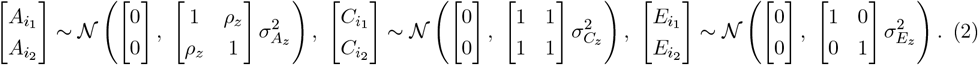

*The relatedness ρ*_*z*_ for the additive genetic effects between two twins *i*_1_ and *i*_2_ in a family is 1 when *z* = MZ and 0 when *z* = DZ. As a side note, the main notations used in this paper are listed in Table 1.

**Table 1:**
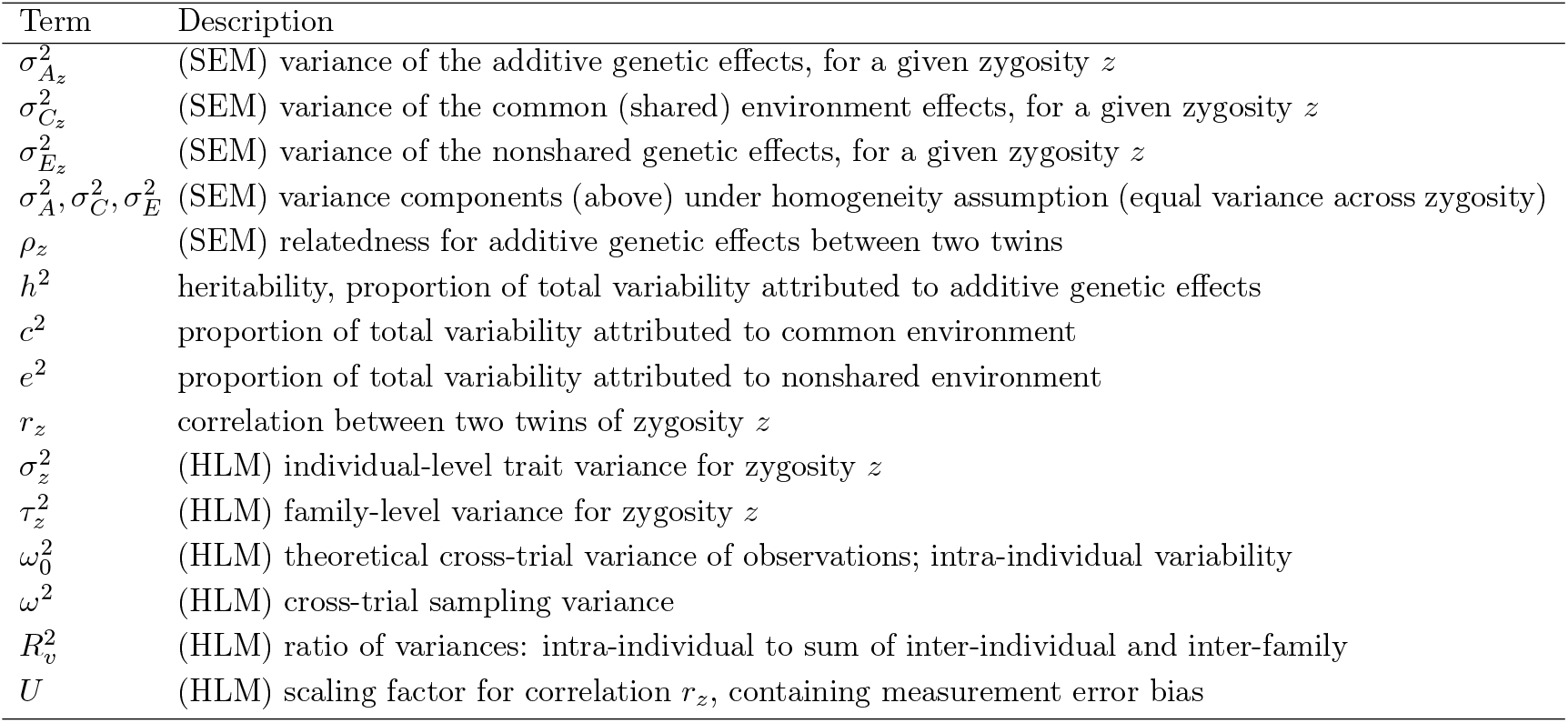
A reference table of major variables and parameters used with heritability modeling. Quantities which originate in hierarchical linear modeling (HLM) and structural equation modeling (SEM) are noted explicitly.

| Term | Description |
| --- | --- |
| $\sigma_{A_z}^2$ | (SEM) variance of the additive genetic effects, for a given zygosity $z$ |
| $\sigma_{C_z}^2$ | (SEM) variance of the common (shared) environment effects, for a given zygosity $z$ |
| $\sigma_{E_z}^2$ | (SEM) variance of the nonshared genetic effects, for a given zygosity $z$ |
| $\sigma_A^2, \sigma_C^2, \sigma_E^2$ | (SEM) variance components (above) under homogeneity assumption (equal variance across zygosity) |
| $\rho_z$ | (SEM) relatedness for additive genetic effects between two twins |
| $h^2$ | heritability, proportion of total variability attributed to additive genetic effects |
| $c^2$ | proportion of total variability attributed to common environment |
| $e^2$ | proportion of total variability attributed to nonshared environment |
| $r_z$ | correlation between two twins of zygosity $z$ |
| $\sigma_z^2$ | (HLM) individual-level trait variance for zygosity $z$ |
| $\tau_z^2$ | (HLM) family-level variance for zygosity $z$ |
| $\omega_0^2$ | (HLM) theoretical cross-trial variance of observations; intra-individual variability |
| $\omega^2$ | (HLM) cross-trial sampling variance |
| $R_v^2$ | (HLM) ratio of variances: intra-individual to sum of inter-individual and inter-family |
| $U$ | (HLM) scaling factor for correlation $r_z$, containing measurement error bias |

A crucial feature of the formulation (2) is the inclusion of the *homogeneity assumption*. This assumption is necessary when estimating variances within the model framework in (2). Six variances, namely 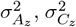, and 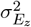 for *z* = DZ and MZ, need to be estimated, resulting in an undetermined system. To resolve this identifiability issue, the variances are assumed to be homogeneous across MZ and DZ twins (Arbet et al., 2020):

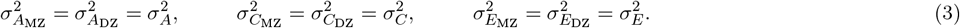

The assumption effectively reduces the number of variance parameters by half. However, this also leads to having homogeneity of total variance between MZ and DZ: 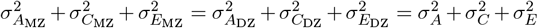

The heritability in a twin study can be defined under the homogeneity assumption (3). The classical methodology is to adopt structural equation modeling (SEM) (e.g., Rijsdijk and Sham, 2002; Holst et al., 2016; Neale et al., 2016; Bates et al., 2019) to estimate the three variances. Then, the three proportions of total variability attributed to additive genetic effects, common environment, and nonshared environment are expressed respectively as,

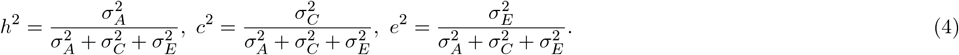

Effect decomposition under the SEM (2) and the associated estimation of heritability can be visually represented as a path diagram (Fig. 1), which is analogous to a directed acyclic diagram in causal inference. The values of *h*^2^, *c*^2^, and *e*^2^ in common practice are typically reported with their point estimates, accompanied by their uncertainty expressed through standard errors or 95% uncertainty intervals^1^.

**Figure 1.**
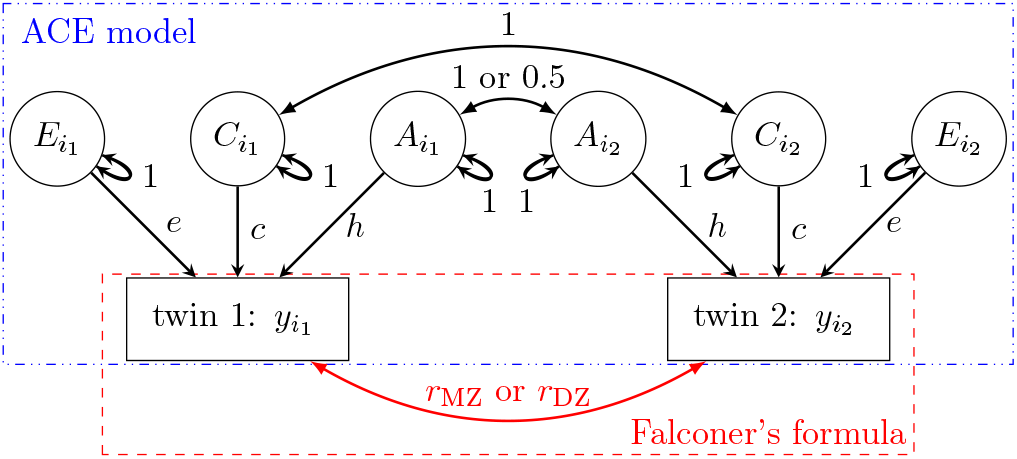
Path diagram (reticular action model) for the ACE formulation *y*_*i*_ = *α* + *A*_*i*_ + *C*_*i*_ + *E*_*i*_. Subscript indices for family (*f*), and zygosity (*z*) are dropped for brevity. Observable effects of *y* are represented as rectangles while latent effects (*A, C* and *E* components) are represented by ellipses. A directed path, represented by a single-headed arrow, indicates predictability. An undirected path, represented by a double-headed curved arrow, indicates a covarying relationship. The value on each path shows the correlation coefficient. Note that *h*^2^, *c*^2^, and *e*^2^ can be estimated through effect partitioning into the three latent components (*A, C*, and *E*), or through Falconer’s formula *h*^2^ = 2(*r*_MZ_ − *r*_DZ_).

The three variability proportions of *h*^2^, *c*^2^, and *e*^2^ can alternatively be expressed as the relatedness between the twins of each zygosity. We denote *r*_MZ_ and *r*_DZ_ as the correlations between two twins *i*_1_ and *i*_2_ within a family *f* for MZ and DZ, respectively. The following can be derived per the ACE model under the SEM formulation (2),

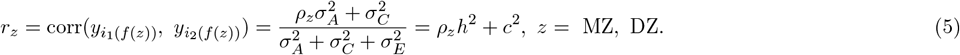

The expression (5) leads to Falconer’s formula (Falconer and MacKay, 1996):

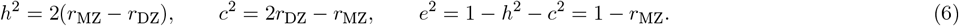

### 1.2 Motivation: addressing the limitations of the conventional SEM framework

We address the challenges of heritability estimation from the perspectives of *accuracy* and *precision*, which are typically the objectives of studies to maximize. Here, we define the accuracy of an estimate as the absence of systematic biases. Inaccuracy, for example, implies an upward or downward shift in a point estimate of effect centrality (e.g., mean, mode, median), an uncertainty interval, or a posterior distribution. On the other hand, we define imprecision as the extent of uncertainty in an estimate, which can be quantified as the standard error or an uncertainty interval.

An implicit assumption is present within the conventional SEM regarding measurement errors, the random or stochastic fluctuations in a measurement from one instance to another. Specifically, the SEM formulation assumes one of two possibilities: 1) the absence of measurement errors in the phenotypic data *y*_*i*(*f* (*z*))_, or 2) the inclusion of measurement errors within the nonshared environment component *E*_*i*(*f* (*z*))_ (along with its associated variance 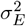) when they are present. Both assumptions lead to the same practical outcome: data with repeated measures are typically preprocessed by aggregating information through a centrality metric (such as the average) before estimating heritability.

Measurement errors are not a significant concern for many measures, such as physical traits that can typically be assessed with high precision. We focus here on heritability estimation in situations where measurement errors are not negligible. For example, in a Stroop task where a large number of trials (e.g., 100) are presented in the experiment for each congruent and incongruent condition. To utilize the SEM formulation directly, the data are typically aggregated, and a centrality measure is used as input.

Data aggregation is a common practice in heritability estimation. Examples can be found in the fields of psychometrics (e.g., Smith et al., 2023; Rea-Sandin et al., 2023; Hung et al., 2023; Gustavson et al., 2023; Vellani et al., 2022; Viktorsson et al., 2022; Yeom et al., 2022; Wang et al., 2020; Routledge et al., 2018; Stins et al., 2005; Schachar et al., 2010; Fan et al., 2001) and neuroimaging (Kastrati et al., 2022; van Drunen et al., 2021; Chen et al., 2019; Harper et al., 2019; van der Meulen et al., 2018; Anokhin et al., 2017; Blokland et al., 2011; Matthews et al., 2007; Polk et al., 2007). However, data aggregation can be a double-edged sword, as valuable information can be lost. The impact of ignoring intra-individual variability has been explored in different contexts. For instance, the issue can be traced back to Spearman (1904) who pointed out the underestimation problem in estimating the correlation between two variables when measurement errors occur. It has also been recently shown that, without proper accountability, test-retest reliability can be substantially underestimated (Rouder and Haaf, 2019; Haines et al., 2020; Chen et al., 2021). Even in heritability estimation, underestimation has been revealed in item response theory due to the adoption of the aggregation process in the form of sum–scores (van den Berg et al., 2007; Schwabe et al., 2019). Recently, the bias problem due to measurement errors has been investigated regarding the reliability of polygenic score and that of a phenotypic trait framed under SEM (Pingault et al., 2022).

Here, we employ a hierarchical linear modeling (HLM) approach to account for intra-individual variability. We propose that *measurement errors should not be considered part of the nonshared environment component*, but rather partitioned appropriately within the model hierarchy. In addition, we employ the HLM approach to reexamine the conventional SEM framework, with the latter being a special case of the former. Specifically, we demonstrate that the common practice of data aggregation fails to adequately address the impact of intraindividual variability. With simulations and real datasets, we address the following questions:

1) Does disregarding intra-individual variability result in biased estimation? If so, to what extent?
2) To what degree does intra-individual variability contribute to precision in heritability estimation?
2) Are typical twin study sample sizes sufficient to achieve a proper precision of heritability estimation?

It is important to note that SEM has been extended to accommodate complex hierarchical data structures (e.g., Mehta and Neale, 2005). Although the SEM framework could potentially be applied to our work, we have opted for HLM due to our preference and familiarity. For clarity, our model comparisons are based on the conventional SEM approach for heritability estimation, not on SEM’s extended capabilities.

## 2 Estimating heritability under hierarchical framework

A hierarchical model partitions data variability by mapping the stratified structure to effects at various levels. The modeling approach is well-established in behavior genetics. For example, van den Oord (2001) proposed using HLM to characterize latent genetic and environmental components of variance in extended families. Guo and Wang (2002) derived heritability estimation through direct variance decomposition. McArdle and Prescott (2005) demonstrated the equivalence between conventional SEM and the variance decomposition approach. Hunter (2021) extended this approach to multiple phenotypes, exploring the feasibility of handling nested data and repeated measures. However, the literature lacks a discussion on how to incorporate intra-individual variability.

### 2.1 HLM: reformulating the SEM approach

We construct the following model by decomposing the data *y*_*i*(*f* (*z*))_ into effects through the three indices of individual (*i* = 1, 2, …, *I*), family (*f* = 1, 2, …, *F*), and zygosity (*z* = MZ, DZ):

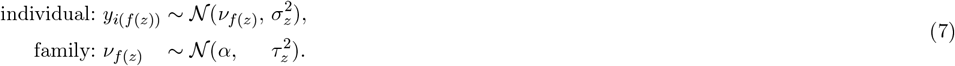

Similar to formula (5), the correlation between any two twins, *i*_1_ and *i*_2_, within a family *f* can be estimated as

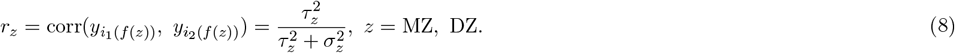

The HLM formulation (7) has been proposed and explored in previous studies in contexts where intraindividual variability is absent (e.g., van den Oord (2001); Guo and Wang, 2002; McArdle and Prescott, 2005). Here, we aim to extend the framework to accommodate cases where intra-individual variability is present. We highlight three advantages regarding the HLM framework. First, the conventional SEM formulation (2) assumes the homogeneity of variances (3), which leads to total variance homogeneity between the two zygosities,

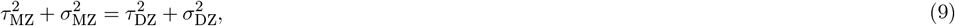

The equivalence between the two modeling approaches can be established when we equate the total variance of 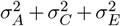 in the SEM formulation (2) and that in (9) under the HLM framework (7), as well as *r*_*z*_ in (5) and (8), leading to:

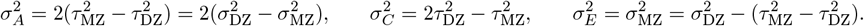

The HLM (7) does not assume variance homogeneity. Instead, a less stringent assumption – proportionality homogeneity across zygosities – is made when using Falconer’s formula (6): the variance proportions accounted for by genetic and shared environmental effects – *h*_*z*_ and *c*_*z*_ (*z* = MZ, DZ) – are the same between the the two zygosities (Arbet et al., 2020). Specifically, this proportionality assumption hinges on the derivation (5) for the Falconer’s formula. With 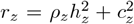, the proportionality assumption of 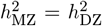 and 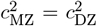 is sufficient for the validity of the Falconer’s formula.

Second, the HLM formulation has the flexibility to accommodate different distribution types. As shown in the formulation (7), the individual-and family-level variances in the two Gaussian distributions 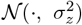 and 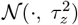 can be expanded to include a wider range of distribution options, such as Student’s *t* and exponentially-modified Gaussian distribution. This flexibility in choosing distributions can greatly enhance the quality of the model, especially when working with datasets that exhibit heavy tails, positive-only quantities, truncated or bounded values. Finally, HLM has the ability to explicitly capture intra-individual variability, rather than grouping measurement errors with the nonshared environment component. It allows for appropriately partitioning variability within the data hierarchy.

### 2.2 Consistency between SEM and HLM

We utilized a publicly available dataset of body mass index (BMI) from the `R` package `mets` (Holst et al., 2016) to validate the HLM approach. Despite having only one BMI measurement per individual, the presence of intra-individual variability can be considered negligible. In summary, the BMI data comprised *I* = 11188 twins from *F* = 6917 families in Finland, including 3665 MZ and 7523 DZ twins aged between 32 and 61 years. Information on each individual’s age and sex was also included in the dataset. Both the data and the code for this example are available at https://github.com/afni/apaper_heritability.

Heritability estimation for the BMI dataset was performed using SEM with the following specifications. Alongside the three latent components *A, C*, and *E* in formulation (1), we incorporated the following covariates: zygosity and a nonlinear age effect for sex using third-order B-spline bases. The SEM formulation was implemented using the `R` packages `mets` and `umx` (Bates et al., 2019), yielding *h*^2^ = 64%, *c*^2^ = 0% and *e*^2^ = 34%.

For the HLM approach, we adopted the model (7) with log-normal and Gaussian density for the individual-and family-level distributions, respectively based on the tendency of the BMI data skewed to the right (Fig. 2A). The following covariates were included: zygosity and nonlinear age effect for each sex using smooth splines with thin plate bases. The model was implemented under the Bayesian framework using the `R` package `brms` (Bürkner, 2017). The resulting *h*^2^, *c*^2^, and *e*^2^ were largely consistent with the SEM estimation (Fig. 2B), both in terms of point estimate and uncertainty range values.

**Figure 2.**
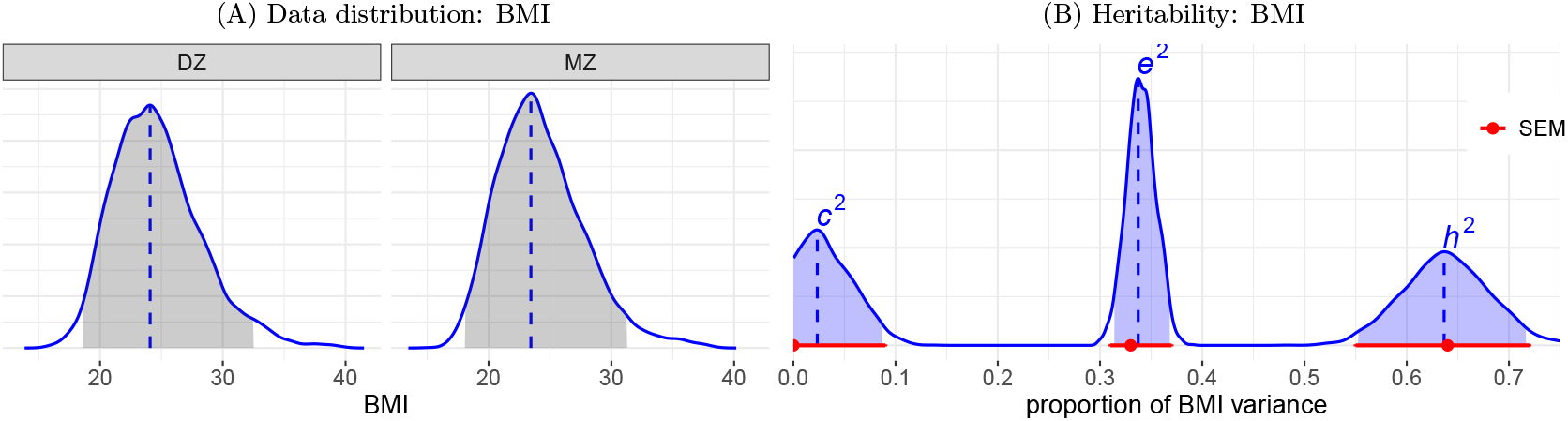
(A) Data distribution. BMI exhibits a right-skewed distribution, with slightly greater dispersion among DZ twins compared to MZ twins. (B) Heritability estimation of BMI data. The proportions of BMI variability attributed to each of the three components are depicted. The distributions for *h*^2^, *c*^2^, and *e*^2^ are estimated using the HLM formulation (7) and presented in blue. The shaded area under each distribution represents the 95% highest density interval, while the vertical dashed line indicates the mode (peak). For comparison, the point estimates (dot) and their corresponding 95% uncertainty intervals (horizontal line) for the SEM (2) are displayed in red.

## 3 HLM: accounting for intra-individual variability

Within the hierarchical framework, we will employ simulations to systematically investigate the influence of intra-individual variability, trial and participant sample sizes on precision, and determine the necessary sample sizes to achieve a satisfactory level of precision. The insights obtained from these simulations will be further validated by applying the HLM framework to a behavioral dataset.

### 3.1 Measurement errors: part of nonshared environment component?

Measurement errors have traditionally been regarded as part of the nonshared environment component in the heritability model, either implicitly or explicitly (e.g., Maes, 2005; Germine et al., 2015). In other words, it has been considered appropriate to include any measurement errors in the trait measurement *y*_*i*(*f* (*z*))_ within the variance component 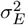 under the SEM (2).

We contend that an ideal modeling approach should appropriately allocate measurement errors rather than grouping them together with the nonshared environment. Suppose that the observation *y*_*i*(*f* (*z*))*t*_ (*t* = 1, 2, …, *T*) in the *t*th trial is sampled from a Gaussian distribution with a mean effect *θ*_*i*(*f* (*z*))_ and a standard deviation *ω*_0_,

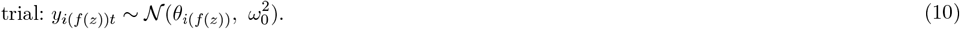

Thus, the sample mean 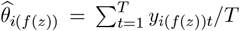 carries a cross-trial sampling variance 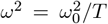, which represents the precision of the estimate. In common practice, when data is aggregated, only the sample mean 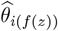 is utilized in the SEM formulation, while the cross-trial sampling variance *ω*^2^ is not explicitly accounted for. Consequently, 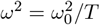 remains embedded as part of the nonshared environment component 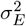, and the estimation of heritability in (4) becomes dependent on trial sample size and sampling precision. As heritability aims to measure differences among individuals rather than within individuals, it is more conceptually sensible to construct a model where the sample size impacts the precision of the estimated variance, rather than its accuracy.

Measurement errors can be appropriately accounted for as a separate component from the nonshared environment. Suppose we consider the individual-level trait effect *θ*_*i*(*f* (*z*))_ as a latent variable. For the corresponding estimate 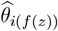, we characterize the measurement errors through the cross-trial sampling variance *ω*^2^. In other words, we do not solely partition the trait into the three latent components of *A, C*, and *E*. Instead, we also treat the true trait effect *θ*_*i*(*f* (*z*))_ as another latent effect, as depicted in the path diagram shown in Figure 3. Additionally, we emphasize that the cross-trial sampling variance *ω*^2^ is not conceptualized as part of the (latent) nonshared environment component *E*, but rather as a separate entity that is directly observable. Therefore, the original SEM (2) is augmented to include two levels,

**Figure 3.**
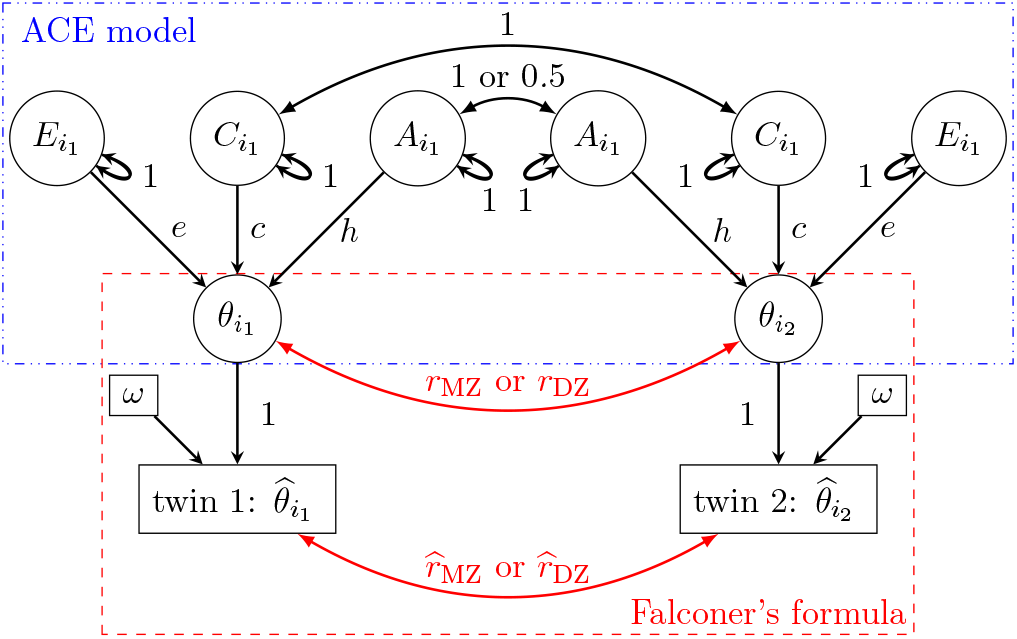
Path diagram (reticular action model) for the augmented SEM (11) and the HLM (12). Subscript indices for family *f*, and zygosity *z* are omitted from the nodes for brevity. Unlike the path diagram in Fig. 1, the trait sample means across trials, 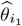 and 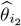, are observable. A directed path, indicated by a single-headed arrow, represents predictability, while an undirected path, shown by a double-headed curved arrow, represents a covarying relationship. The value on each path indicates the correlation coefficient.

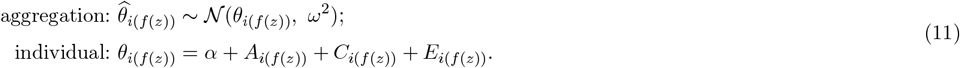

The same distribution assumptions in (2) apply to the three latent effects of *A, C*, and *E* here. Solving this augmented SEM (11) directly is challenging. However, in Subsection 3.4, we will present an approximate approach to heritability estimation.

Measurement errors can also be incorporated into the HLM framework. In particular, we augment the HLM formulation (7) to

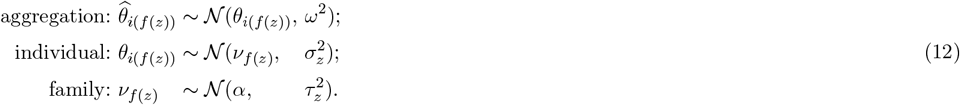

When the cross-trial sampling variance *ω*^2^ is available, this augmented model (12) can be analyzed under the Bayesian framework or approximately solved as discussed in Subsection 3.4.

Framing the presence of measurement errors as a distinction between observed and latent effects helps in understanding the associated impact. As illustrated in the path diagram (Fig. 3), one can estimate heritability directly based on the correlations 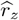 using the sample means 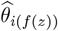 through Falconer’s formula (6). Long ago, Spearman (1904) highlighted a bias issue: the correlation between two quantities becomes attenuated when measurement errors are not accounted for. Similarly, when measurement errors are disregarded, 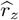 would be underestimated compared to their counterparts *r*_*z*_ based on the latent effects *θ*_*i*(*f* (*z*))_ (Fig. 3). Next, we construct an HLM formulation at the observation level to fully illustrate the issues associated with data aggregation.

### 3.2 HLM with trial-level data under one task condition

We begin by extending the HLM (7) to a case where data are collected at the observation level with repeated measures through trials under a single task condition. The data *y*_*i*(*f* (*z*))*t*_ are represented using four indices: family (*f* = 1, 2, …, *F*), zygosity (*z* = MZ, DZ), individual (*i* = 1, 2, …, *I*), and trial (*t* = 1, 2, …, *T*). Instead of utilizing aggregated information as in (11) and (12), we formulate the following HLM based on (10):

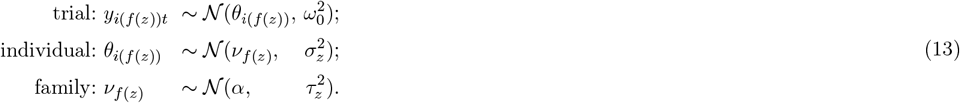

The corresponding path diagram is shown in Fig. 4. The formulation (13) can be extended to include various distributions. For instance, the trial-level effects can be characterized through Bernoulli, gamma, and Poisson distributions, which allow for modeling different types of data (count, binary, skewed).

**Figure 4.**
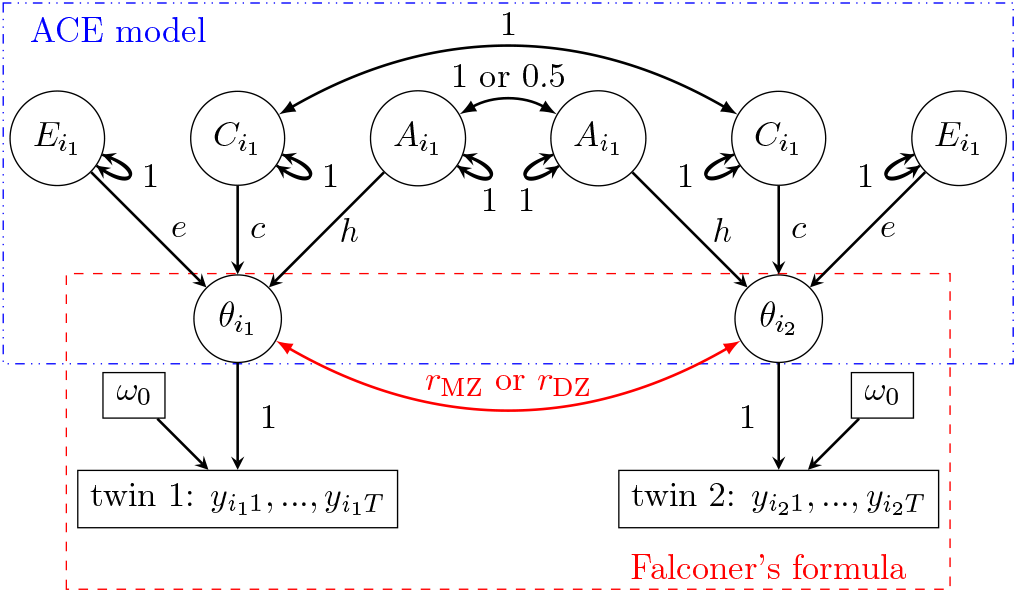
Path diagram (reticular action model) for the HLM formulation (13). Subscript indices for family *f*, and zygosity *z* are omitted from the nodes for brevity. Unlike the path diagram in Fig. 3, the trait measures *y* are observable. A directed path, indicated by a single-headed arrow, represents predictability, while an undirected path, shown by a double-headed curved arrow, represents a covarying relationship. The value on each path indicates the correlation coefficient.

Heritability can be estimated using the HLM (13) for trial-level data as follows. Similar to the situation with HLM for the conventional SEM in (7), we compute the correlations between two twins within a family using the inter-individual and inter-family variances, 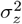 and 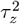, through the formula (8). Then, the three variance proportions *h*^2^, *c*^2^, and *e*^2^ are obtained using Falconer’s formula (6).

### 3.3 Consequences of data aggregation under the SEM formulation

Now we examine the common practice of data aggregation in light of the HLM (13). To capture the crucial role of intra-individual variability 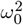 across different phenotypic traits, we define a dimensionless measure of the variability ratio for each zygosity:

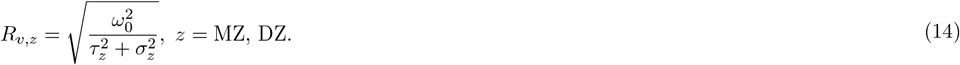

The variability ratio *R*_*v*,*z*_ captures the fundamental aspect of heritability: the proportion of inter-family and inter-individual variance relative to intra-individual variance. Under the assumption of homogeneity (9), *R*_*v*,MZ_ = *R*_*v*,DZ_. For simplicity, we drop the subscript *Z* and denote their average as *R*_*v*_. When the trial-level data *y*_*i*(*f* (*z*))*t*_ are aggregated across trials with their average 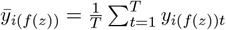, the model (7) becomes

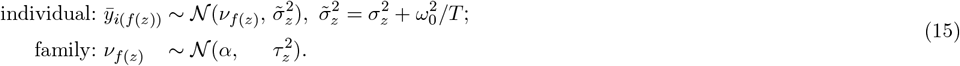

In comparing the model (15) with aggregated data to its counterpart (7) for data without measurement errors, we note that ignoring intra-individual variability leads to its combination with the inter-individual variance 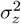 into 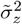. As a result, the correlations *r*_MZ_ and *r*_DZ_ in (8) are updated to

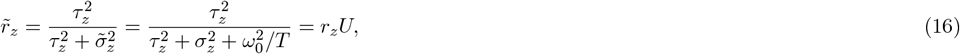

where the introduced bias into *r*_MZ_ and *r*_DZ_ is quantified by the dimensionless quantity 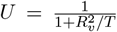. It is noteworthy that when *ω*_0_ is nonzero, *U <* 1, signifying that 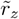 is consistently downward biased. This bias arises due to the presence of intra-individual variability, and its attenuation rate follows a sigmoid function of *R*_*v*_. In the limit where 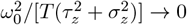, which can occur with decreasing standard error *ω*_0_ or increasing trial size *T*, 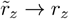.

The parameter *U* quantifies the degree of bias in heritability estimation under SEM when data aggregation is applied. According to Falconer’s formula (6), both *h*^2^ and *c*^2^ would be underestimated by a factor of *U*, while *e*^2^ would be overestimated by the same factor. For example, with *T* = 100 trials, a small intra-individual variability such as *R*_*v*_ = 1 has negligible impact on heritability estimation (*U* ≈ 0.99), whereas a large intra-individual variability with *R*_*v*_ = 10 substantially underestimates *h*^2^ and *c*^2^ by 50%. Conversely, if *R*_*v*_ = 3, biases cannot be disregarded even with *T* = 20 trials unless *T* approaches or exceeds 100.

One direct way to view the distinction between SEM and HLM is to compare their respective path diagrams (Figs. 3 and 4). HLM preserves the hierarchical structure and cross-trial variability, ensuring this information propagates across other hierarchical levels (Eq. (13)). In contrast, SEM obscures this variability through data aggregation, compromising the hierarchical integrity. This loss of data structure fidelity in SEM leads to biased underestimation of heritability, as captured through the parameter *U* in the expression (16).

### 3.4 Ameliorating the biases in the SEM formulation

The biases induced in the SEM formulation can be theoretically corrected by introducing an adjustment term in the denominator of (16) to counteract the contaminating effect of 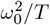 under the model (15),

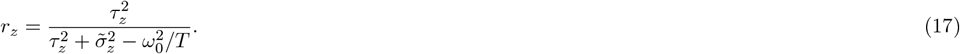

Similarly, decontamination can be achieved for the formulas (4) and (8). However, these adjustments rely on the availability of the intra-individual variance 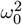, which is not directly accessible once the data are aggregated.

Nevertheless, the biases can be practically mitigated. For example, we can use the cross-trial variance estimates 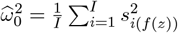 in (17), leading to the following approximate adjustment,

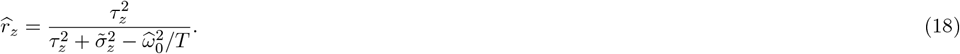

Similarly, as *σ*_*E*_ inherently contains the additive contribution of measurement error, we can directly adjust the biases in the SEM estimates to,

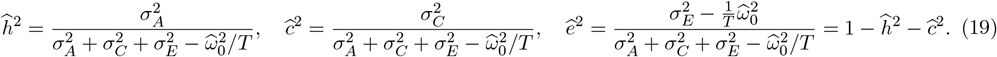

These approximate adjustments (18) and (19) offer a solution to the augmented SEM formulation (11) and its hierarchical counterpart (12). We will further explore and validate their effectiveness as approximate adjustments later with an experimental dataset.

An intriguing aspect of biased estimation for heritability is its analogy to the phenomenon of correlation attenuation in the presence of measurement errors. Spearman (1904) recognized the problem of bias caused by measurement errors and proposed an adjustment method to disattenuate the correlation between two variables. In (16), the term *U* serves a similar purpose to the reliability coefficient or separation index in classical test theory. Consequently, it is interesting to note that the decontamination formula (18) and its approximation (19) employ a similar adjustment strategy as suggested by Spearman (1904).

### 3.5 HLM with trial-level data under two task conditions

We now extend the HLM (13) to accommodate two task conditions. In fields such as psychometrics and neuroimaging, the focus often lies in comparing and analyzing the contrast between two conditions. We expand the previous HLM (13) to one with hierarchical levels using five indices: family (*f* = 1, 2, …, *F*), zygosity (*z* = MZ, DZ), individual (*i* = 1, 2, …, *I*), condition (*c* = *c*_1_, *c*_2_), and trial (*t* = 1, 2, …, *T*):

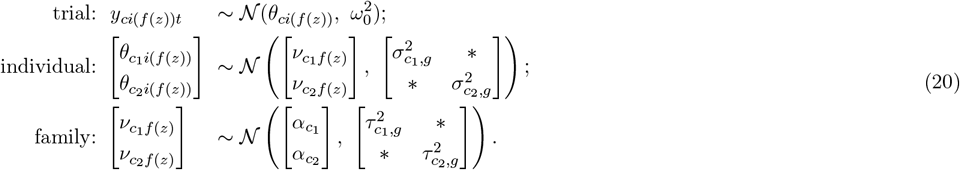

The differences from subsection 2.2 are twofold: (a) the presence of two intercepts, *α*_*c*_, one for each condition, and (b) the individual-and family-level distributions being bivariate instead of univariate. While the covariances for the individual-and family-level distributions are not of interest in the current context, we acknowledge their presence by denoting them with an asterisk * in the respective variance-covariance matrix.

Heritability estimation is straightforward for each condition under the HLM (20). First, we calculate the correlation between two twins within a family for condition *c*_*k*_ (*k* = 1, 2) using

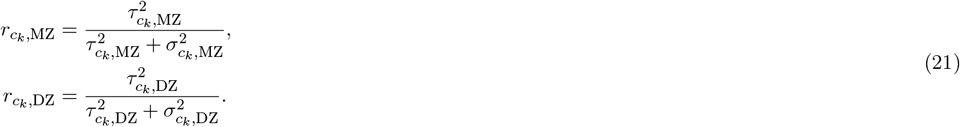

Then, we estimate heritability for each condition by plugging 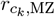 and 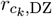 into Falconer’s formula (6).

Two approaches are available for estimating heritability and the variability ratio for the contrast between two conditions. One approach involves reparameterizing the model (20) through an indicator variable using effect coding for the two conditions,

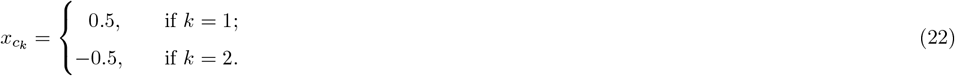

Alternatively, within the Bayesian framework, one can directly obtain the distribution of the contrast by formulating it based on each condition’s posterior draws from the HLM (20).

Biased estimation due to data aggregation and its adjustment in the previous subsection also apply to the case with two conditions. For each condition and their contrast, the variability ratio *R*_*v*_ can be similarly defined.

The only modification for the contrast is to replace 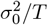 with 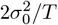 in (14). Similarly, the discussions regarding the biases in heritability estimation for each condition in the preceding subsection can be directly applied here.

However, when considering the contrast, the bias requires replacing 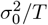 in (15-19) with 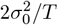.

### 3.6 Assessing model performance through simulations

We have shown that, by closely reflecting the actual data structure, an HLM framework appropriately accounts for different sources of data variability. However, the precision of heritability estimation remains unclear, as there is no analytical quantification available. Simulations were conducted to assess the precision of heritability estimation from the following aspects regarding the impact of intra-individual variability that the analytical approach cannot easily reveal: 1) the precision of heritability estimation, and 2) the requirement of family and trial sample sizes in twin studies.

Detailed information about the simulations and results can be found in Appendix A. As expected, HLM showed no bias in heritability estimation, whereas biases under the SEM framework become more pronounced as the variability ratio *R*_*v*_ increases and/or the trial sample size *T* decreases. Additionally, biases under the SEM formulation with aggregated data can be effectively adjusted using empirical or theoretical standard errors through (17) or (18). More importantly, simulations indicate that intra-individual variability impacts estimation precision. Specifically, larger *R*_*v*_ leads to poorer precision. Lastly, family sample size has a greater impact than trial sample size on estimation precision. For instance, when *R*_*v*_ ≲ 1, an appropriate level of uncertainty can be achieved with 50 trials and 1,000 families. When *R*_*v*_ ≫ 1, several thousand families may be necessary.

## 4 Applying HLM to an experimental dataset

We apply the HLM approach to an experimental dataset to address two primary questions. First, do the insights gained from numerical simulations in the previous section align with the findings when real data is analyzed? Second, what is the range of the relative magnitude of intra-individual variability, as indicated by the ratio *R*_*v*_, in commonly encountered empirical datasets?

We utilize a behavioral dataset obtained from an experiment conducted as part of the ABCD study. The experiment investigated selective attention during adolescence using an emotional Stroop task (Smolker et al., 2022), with reaction time (RT) as a phenotypic trait. The data was collected during the 1-year follow-up visit and is publicly accessible through the 2020 ABCD Annual Curated Data Release 4.0 (https://nda.nih.gov/study.html?id=1299). The analysis scripts can be found online at: https://github.com/afni/apaper_heritability.

### 4.1 Data description

A subset of the original dataset, specifically containing twins, was utilized for the analysis. Refer to Table 2 for detailed information on participant counts and demographic data. The subset consisted of 1,102 twins (including some triplets) from 555 families and was selected from a larger dataset of 11,876 participants (Iacono et al., 2018). Among the included twins, there were 461 MZ and 641 DZ participants.

**Table 2:**
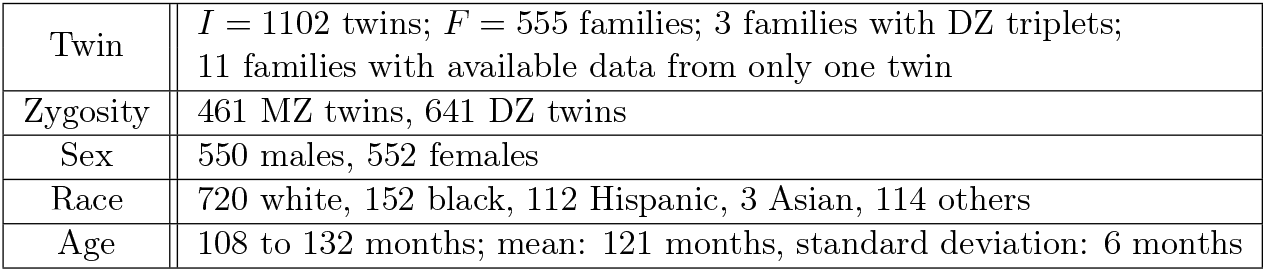
Demographic information of twins in a Stroop experiment from the ABCD Study.

We focus on the RT data for estimating heritability. The RT was measured for two levels of congruency in a Stroop task: congruent and incongruent. Each participant was instructed to respond to a total of 48 trials, consisting of 24 congruent and 24 incongruent trials. The response window for each trial was set to 2000 ms. Among 1,102 twins, a total of 49,524 trials were included in the analysis after excluding incorrect responses. This resulted in an overall correct response rate of approximately 93.6%. The distribution of RTs is heavily right-skewed (Fig. 5), with a mode of 962 ms and a 95% highest density interval ranging from 634 to 1,827 ms.

**Figure 5.**
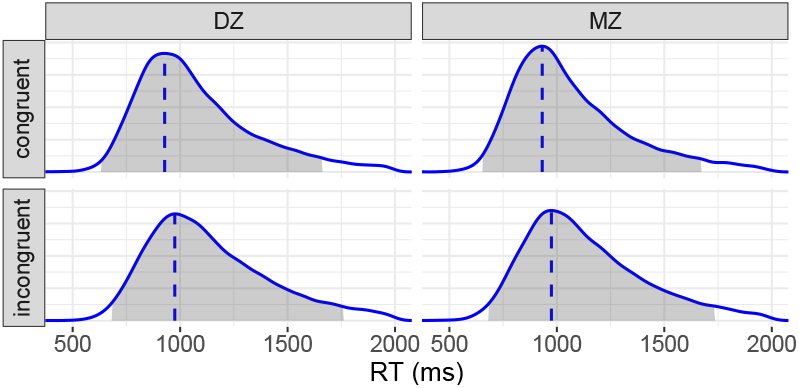
Reaction time (RT) distributions. The shaded area under each density represents the 95% highest density interval, while the vertical dashed line indicates the mode. The RT distribution is 1) right-skewed, 2) slightly right-shifted under the incongruent condition compared to the congruent condition, and 3) slightly more dispersed for DZ twins than MZ twins.

### 4.2 Model comparisons with real data

The RT data was analyzed using two different approaches: HLM and SEM. For HLM, the trial-level data was fitted using the formulation (20). Site, sex, race, age, and zygosity were included as covariates. To account for the right-skewness (Fig. 5), a log-normal distribution was used for the trial-level effects. The Bayesian framework was employed to implement the HLM approach, utilizing the `brms` package in `R` (Bürkner, 2017). Heritabilty was estimated for each of the three RT effects: the congruent condition, incongruent condition, and the Stroop effect (the contrast between incongruent and congruent conditions). The SEM was applied with aggregated data. Specifically, RT was aggregated across trials for each condition at the individual level. Similar to the HLM approach, covariates including site, sex, race, age, and zygosity were included. The SEM formulation was implemented using the R packages of `mets` and `umx`. The computational time for SEM with aggregated data was negligible. In contrast, HLM-based estimation, using Markov chain Monte Carlo simulations, required 3.5 hours with four chains and 12 threads per chain. These computations were performed on an Intel Server S2600WFT equipped with 96 CPUs running at 2933 MHz.

The estimation results are presented in Figure 6. The performance of HLM relative to SEM, as summarized below, aligns with our simulations in subsection 3.6. Overall, both SEM and HLM exhibited a significant amount of uncertainty in estimating heritability for both conditions. SEM displayed noticeable underestimation of *h*^2^ and *c*^2^ (first two columns, Fig. 6). However, neither modeling approach provided satisfactory estimation for the contrast (third column, Fig. 6). This estimation challenge arises from the combination of three factors encapsulated by a large intra-individual variability ratio *R*_*v*_ ≈ 10: 1) a much smaller effect size, 2) an extremely limited trial sample size, and 3) a relatively small twin sample size.

**Figure 6.**
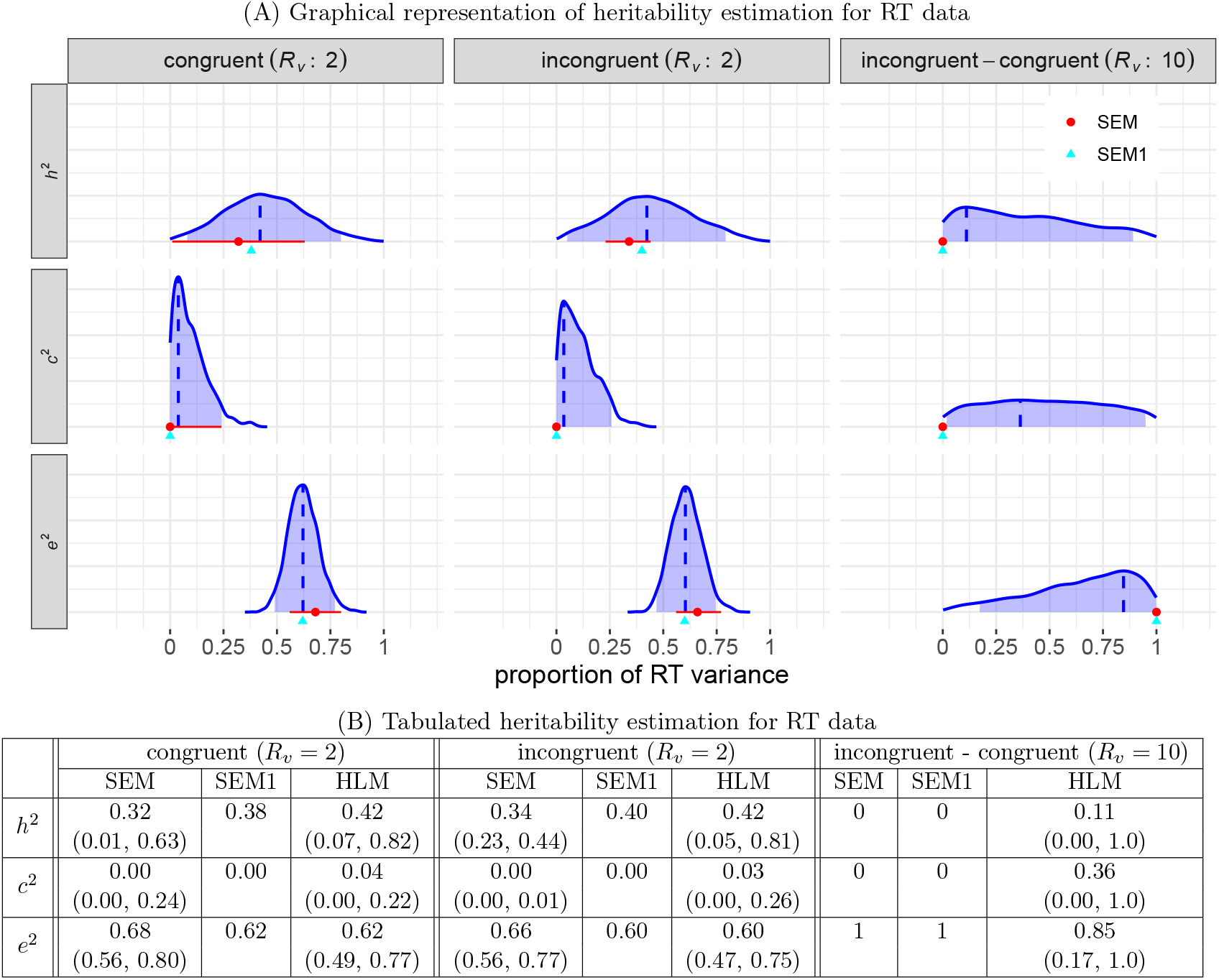
Estimated heritability for RT data. (A) The three columns represent the two conditions (congruent and incongruent), as well as their contrast, with the corresponding variability ratio *R*_*v*_ indicated in the column labels. The three rows correspond to *h*^2^, *c*^2^, and *e*^2^. In each panel, the HLM result is represented by a solid blue density curve, derived from random draws from posterior chains. The mode is marked by a vertical blue dashed line, and the shaded blue region represents the 95% uncertainty interval. The SEM counterparts are also displayed in each plot, with the point estimate depicted as a red dot and its 95% uncertainty interval represented by a horizontal red line. Notably, the SEM point estimates tend to be smaller than their HLM counterparts for *h*^2^ and *c*^2^ (while larger for *e*^2^). Adjustments for the SEM estimates, using the formula (19), are denoted as SEM1 and shown as green triangles. (B) Comparisons among the three models are presented with their point estimates and 95% uncertainty intervals. For the HLM, estimates are derived from the modes and highest density intervals of the posterior distributions in (A).

Below are a few detailed elaborations:

1) **Impact of relative intra-individual variability on estimation precision**. First, under each of the individual congruent and incongruent conditions, the observed intra-individual variability was not large (*R*_*v*_ ≈ 2), resulting in moderate uncertainties for HLM estimates of *h*^2^, *c*^2^, and *e*^2^. However, the SEM approach showed difficulty in accurately assessing uncertainty near the parameter boundaries (e.g., 0 or 1 for *h*^2^, *c*^2^, and *e*^2^). For instance, the SEM’s small uncertainty interval (0, 0.01) for *c*^2^ likely stemmed from numerical singularity issues in the traditional statistical framework. In contrast, regularization in hierarchical modeling (Chung et al., 2013) yielded a more reasonable uncertainty interval (0, 0.28) for *c*^2^ under the Bayesian framework. Second, the intra-individual variability for the contrast between the two conditions was large (*R*_*v*_ ≈ 10). Consequently, the uncertainties for *h*^2^, *c*^2^ and *e*^2^ were very large, with the estimated density of *c*^2^ resembling a uniform distribution. The SEM approach did not provide any meaningful estimates either, and this challenge was further demonstrated by its inability to provide an appropriate uncertainty interval, yielding only a single point estimate constrained at the parameter boundaries, likely due to convergence difficulties.
2) **Impact of relative intra-individual variability on estimation bias**. Under both conditions, the intra-individual variability is moderate (*R*_*v*_ ≈ 2), resulting in small underestimations of *h*^2^ and *c*^2^ by SEM. The overestimation of *e*^2^ was also small. However, the large intra-individual variability for the contrast (*R*_*v*_ ≈ 10) led to more noticeable underestimations of *h*^2^ and *c*^2^ by SEM.
3) **Impact of relative intra-individual variability on sample sizes**. The larger *R*_*v*_ for the contrast (*R*_*v*_ ≈ 10) is consequential. Simulation results in subsection 3.6 indicate that larger sample sizes, especially in terms of family count, would be required to reduce the large uncertainty. We note that the observed range of *R*_*v*_ values aligns with psychometric data from individual studies in test-retest reliability estimation (Rouder and Haaf, 2019; Chen et al., 2021; Baker et al., 2021).
4) **Bias adjustment for SEM estimates**. The biases under the SEM framework, due to data aggregation, adjusted using formula (18), were reduced. The adjusted estimates for 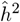 under the congruent, incongruent, and contrast conditions were 0.38, 0.40, and 0.0, respectively (green triangles, Fig. 6). These adjustments effectively reduced bias, although they remained slightly biased compared to HLM, which could be attributed to deviations from the Gaussian assumption under SEM.

We also explored the HLM approach for the ABCD-Stroop data using a conventional linear mixed-effects modeling framework instead of a Bayesian approach. The `lmer` function from the `lme4` package in R (Bates et al., 2015) was utilized to fit the models (13) and (20) with RT log-transformed. Although the point estimates (not shown here) for *h*^2^, *c*^2^, *e*^2^, and *R*_*v*_ under the congruent and incongruent task conditions were largely consistent with the values obtained under the Bayesian framework, the numerical solver in `lme4` failed to converge for the contrast between the incongruent and congruent conditions due to the relatively small inter-individual variances (*R*_*v*_ ≈ 10).

## 5 Discussion

Heritability estimation based on data with a non-negligible intra-individual variability necessitates a model that accurately represents the underlying hierarchical structure. When assessing a phenotypic trait with repeated measures, we propose a hierarchical model that encompasses all relevant levels, allowing for the incorporation, propagation, and separation of intra-individual measurement error from parameter estimation at higher levels. Through numerical simulations and a real behavioral dataset, we have demonstrated a few advantages of HLM. These advantages include: 1) avoidance of estimation bias, 2) the ability to account for the significant influence of intra-individual variability on heritability estimation, and 3) enhanced interpretability and explanatory power of results, such as identifying the challenges associated with reducing estimation uncertainty due to sample size limitations.

### 5.1 The importance of modeling data generating mechanism

Data reduction through aggregation is a commonly used in statistical applications, particularly in studying individual differences. Even though intraindividual variability has long been recognized (Fiske and Rice, 1955), many classical frameworks have been applied in contexts where measurement errors are minimal or nonexistent. For example, intraclass correlation (Fisher, 1954) for test-retest reliability in individual differences is typically assessed without considering intra-individual variability. A similar situation can be observed in heritability estimation: to date, common modeling approaches do not explicitly account for the level of measurement units (e.g., individual trials), and instead they simplify the data through preliminary aggregation steps such as averaging.

Can data aggregation be justified by attributing intraindividual variability to the nonshared environment (the *E* component in the ACE model)? The underlying rationale for data aggregation is that intraindividual variability arises either from true biological fluctuations (ontological variability) or measurement limitations (epistemological noise). However, the specific sources are often too complex to be fully accounted for in typical studies. A more pragmatic and effective approach is to adopt a causal inference perspective that focuses on the underlying data-generating process. It is well established that when intraindividual variability follows systematic patterns, treating it solely as residual errors can introduce bias and misinterpretation. Instead, explicitly modeling this variability is crucial to ensuring accurate and meaningful estimates.

In heritability estimation, path diagrams are commonly used to depict causal relationships among variables. Within this framework, a latent trait or condition is conceptualized as a higher-level theoretical construct that causally influences each individual measurement or trial (see Fig. 4). In other words, individual measurements are specific realizations determined by the underlying latent construct. Crucially, heritability is defined at the level of this latent construct, not at the level of single measurements.

A hierarchical modeling framework more accurately maps the causal structure outlined in the path diagram. In contrast, conventional SEM, which treats intraindividual variability as residual errors, does not fully align with the causal structure and can lead to underestimated heritability, as demonstrated in Subsection 3.3. This underscores the necessity of explicitly modeling hierarchical data structures to account for both between-and intraindividual variability.

Our empirical evidence here from both a real dataset and simulations demonstrates that adopting an HLM framework—which respects the full hierarchical structure of the data—yields more accurate heritability estimates than the conventional SEM for psychometric traits. When intraindividual variability is minimal—as is often the case with many physical traits—the conventional SEM can be seen as a special asymptotic case of a more general HLM, and aggregation is justified because the impact of such variability is negligible (*R*_*v*_ ≪ 1). However, for traits such as psychometric measures where intraindividual variability is substantial (*R*_*v*_ ≫ 1), simply aggregating data (i.e., incorporating this variability into residual errors) will likely lead to biased heritability estimates. This underestimation issue extends beyond heritability estimation and has also been observed in test-retest reliability estimation (Rouder and Haaf, 2019; Haines et al., 2020; Chen et al., 2021) and in neuroimaging experimental designs, where the role of trial samples is often overlooked (Chen et al., 2022).

To recapitulate, the HLM approach acknowledges that heritability is defined at the latent trait level and avoids biases associated with oversimplified data aggregation. While data aggregation may be acceptable for traits with minimal intraindividual variability, a hierarchical modeling approach that directly incorporates the causal structure of the data is essential for accurately estimating heritability when variability is pronounced.

To improve the accuracy of heritability estimation, we recommend adopting HLM in the presence of intraindividual variability. It is important to note that the HLM framework is not mutually exclusive with SEM, on which SEM and other methods such as common pathway model are based. As the path diagrams in Figs. 1 and 4 illustrate, both frameworks are conceptually consistent, as discussed in subsection 2.2. Nevertheless, we emphasize that the broader framework of HLM combined with the Bayesian approach offers several advantages:

1) It supports a wider range of numerically implemented distributions (e.g., Student’s *t*, inverse Gaussian).
2) It integrates uncertainty assessment into a single process.
3) It robustly handles variance-covariance structures.

While the last point is important for theoretical and interpretational reasons, it also has useful practical benefits. In the commonly-used R software packages, there are severe challenges faced when using methods implemented in the `nlme` and `lme4` packages, which can struggle with numerical singularities when correlations (or variances) approach boundary values such as-1, 1, or 0, as encountered in this Stroop dataset. The proposed framework avoids these difficulties.

### 5.2 Biases, uncertainty and challenge of heritability estimation

There are two aspects of accuracy compromise that need to be considered in heritability estimation. The first aspect pertains to estimation biases. As demonstrated in this study, failure to fully incorporate the data structure can lead to biased estimates of heritability. The second aspect concerns the uncertainty in heritability estimation. In addition to providing a point estimate for the effect of interest, it is equally important to quantify its uncertainty, characterized through measures such as standard error, an uncertainty interval (e.g., 95%), or even a full distribution (as depicted in Fig. 6). However, uncertainty is often not well emphasized in common practice. In some cases, only the central tendency (e.g., mean) of heritability estimation is reported. However, to truly comprehend the generalizability of results, understanding uncertainty is crucial. One of the benefits of the HLM framework is its ability to directly generate posterior distributions that illustrate estimate precision.

In the presence of intra-individual variability, one might be tempted to adopt the bias adjustment approach using the conventional SEM. Our findings demonstrate that the biases resulting from data aggregation can be mitigated to some extent if variability can be determined separately (e.g., through repeated measures), as indicated by formulas (18) or (19). However, in practical applications, these adjustments are suboptimal due to distributional deviations, as demonstrated in our example using the Stroop dataset. Furthermore, an effective adjustment for biases in uncertainty assessment is currently lacking. Hence, a comprehensive HLM framework remains the preferred choice.

Sample sizes remain a challenge in twin studies. The dataset we used for demonstration, exemplifying a cognitive inhibition study, suggests that reasonable levels of uncertainty can be achieved with sample sizes of less than 1000 families and less than 100 trials per individual condition (congruent and incongruent). The estimated heritability of approximately 40% (first two columns, Fig. 6) aligns with the general range observed in typical phenotypic traits in the literature (Polderman et al., 2015). However, the contrast between conditions is often the focal point of interest. Even with HLM estimation, the uncertainty of heritability for this contrast remains unresolved (third column, Fig. 6), creating imprecision regarding its magnitude. In other words, despite attempts by the Consortium (Iacono et al., 2018; Smolker et al., 2022) to address the sample size issue, the dataset from the ABCD Study (consisting of 461 MZ and 641 DZ twin pairs, with less than 48 trials per condition) does not provide sufficient certainty for estimating the heritability of the Stroop effect. Achieving a reasonable level of precision may require impractical sample sizes (e.g., hundreds of trials and thousands of individuals).

The relative magnitude of intra-individual variability, as quantified by the ratio *R*_*v*_, serves as an informative indicator in heritability modeling. As a dimensionless parameter, it influences not only the accuracy and uncertainty of heritability estimation but also those of test-retest reliability (Rouder et al., 2023; Chen et al., 2021). Historical power analyses in twin studies have suggested a minimum sample size of 600 twin pairs (Martin et al., 1978; Sham et al., 2020). However, our simulations demonstrate that, in the presence of measurement errors, a large *R*_*v*_ poses a significant challenge for future studies in the field of individual differences, particularly when examining effect contrasts and higher-order interactions. Additionally, this ratio highlights the relative importance of trial sample size compared to participant sample size across various experimental modalities, such as functional magnetic resonance imaging, magnetoencephalography, electroencephalography, and psychometrics. In all these cases, the *R*_*v*_ ratio exceeds 1, and sometimes even surpasses 10 (Baker et al., 2021; Chen et al., 2021; Chen et al., 2022). Due to this substantial ratio, the sample size of trials can be nearly as crucial as the number of participants in terms of experimental efficiency in neuroimaging and psychometrics.

### 5.3 Heritability estimation in neuroimaging

To date, there has been an increasing number of twin studies utilizing task-based functional magnetic resonance imaging (fMRI). In these studies, it has been a common practice to aggregate data across trials during fMRI data analysis, resulting in the neglect of intra-individual variability, which is neither accounted for nor reported. For instance, Polk et al. (2007) reported a small heritability estimate of blood oxygenation level-dependent (BOLD) response (*h*^2^ ∼ 0.2) for face and house processing (the specific contrasts were not analyzed), but negligible heritability for pseudowords and chairs in the ventral visual cortex, based on an fMRI experiment involving 13 MZ and 11 DZ twins, with 90 trials per condition. Similarly, Matthews et al. (2007) revealed a moderate heritability (*h*^2^ = 0.37; 90% interval: (0, 0.74)) for the interference effect in the dorsal anterior cingulate cortex during a multi-source interference task with congruent and incongruent conditions, involving 20 MZ and 20 DZ twins, with 144 trials per condition. The heritability estimates for other regions were negligible. In contrast, the heritability of reaction time was moderate for the congruent condition (*h*^2^ = 0.45; 90% interval: (0, 0.76)), but negligible for the incongruent condition and the interference effect. Additionally, Blokland et al. (2011) found moderate to high heritability estimates (*h*^2^ = 0.40 to 0.65) in more than ten regions during an n-back working memory experiment involving 150 MZ and 132 DZ twins, with 128 trials per condition.

Is intra-individual variability a concern for heritability estimation in neuroimaging? The aforementioned task-based fMRI experiments have primarily relied on a large number of trials to obtain reliable estimates of condition-level effects. This is similar to the Stroop dataset we investigated here, with the distinction that the focus is on BOLD response rather than reaction time. However, the family sample size has often been relatively small, leading to larger uncertainty ranges in the estimates. This issue is particularly pronounced because the relative intra-individual variability across the brain, as reported in the literature, tends to be substantial, with *R*_*v*_ ≫ 1 (Chen et al., 2021; Baker et al., 2021; Chen et al., 2022). Thus, heritability estimation in neuroimaging is at least as challenging as typical traits such as psychometric data.

### 5.4 Limitations of heritability estimation through HLM

The HLM approach comes with additional costs. Firstly, introducing an extra level in the data hierarchy significantly increases the complexity of the model structure. Secondly, and perhaps the greater challenge, this increased complexity brings along numerical burdens. Traditional tools like linear mixed-effects estimation are not well-suited for solving hierarchical models of heritability. Instead, resorting to a Bayesian approach may be necessary to handle the numerical challenges (e.g., singularity).

There is always room for improvement in modeling. For example, the full details of the underlying mechanism and framework of cognitive inhibition involved in the reaction time of the Stroop effect are not fully known to researchers. Therefore, no model can fully replicate their structure. However, HLM attempts to model as much as is known and observed in a study. Model fitting can be improved by incorporating auxiliary information, such as accommodating abnormalities like skewness, outliers, and truncation through more inclusive and adaptive distributions (e.g., log-normal, ex-Gaussian). Additionally, one could reconsider the chosen partitioning into three components of *h*^2^, *c*^2^, and *e*^2^ in twin studies and other assumptions (Robette et al., 2022): the additivity of genetic effects, the absence of assortative mating, the nonexistence of genetic dominance or epistasis, the generalizability from twins to the rest of the population, equal environment impact between MZ and DZ twins, and the absence of gene-environment correlation or interaction.

Further integrating HLM with the conventional SEM framework presents a promising avenue for future research. SEM, with its long-established history, offers distinct advantages, including intuitive interpretation, specialized applications, and computational efficiency. While beyond the scope of this study, leveraging the strengths of both SEM and HLM (e.g., Mehta and Neale, 2005) holds significant potential. A unified approach could enable greater flexibility in modeling distributions, account for intraindividual variability, and reduce estimation biases. In addition, this study focuses on intraindividual categorical variables, such as task conditions. Future research could extend this framework to intraindividual quantitative variables, particularly in the context of longitudinal data (e.g., Eaves et al., 1986).

The interpretation of heritability is subtle and sometimes controversial. Our focus here is solely on the technical aspects of heritability estimation. Nevertheless, we emphasize caution in its interpretation. As a statistical metric, heritability captures variation and/or correlation rather than causation. Therefore, one must not confuse the extent of phenotype variability with the contribution of genetic factors. The concept of heritability effectively pertains to the population level and cannot be realistically applied to a particular individual. On the other hand, the information provided by heritability lies in its potential predictability. It can probabilistically predict, but not causally determine, the extent of phenotypic variability. A high heritability for a phenotypic trait may warrant further investigation into the underlying complex genetic mechanisms or the etiology of genetic risk factors, such as biomarkers. This perspective highlights the need to complement heritability research of variance partitioning with mechanism elucidation (Downes and Turkheimer, 2022).

## 6 Conclusions

We propose an HLM approach to improve heritability estimation in twin studies when the phenotypic trait is measured with multiple samples. The methodology aims to separate measurement errors from the variations of interest and addresses issues such as information loss due to data reduction, distribution violations, and uncertainty characterization in current modeling approaches. We demonstrated that the conventional SEM is likely to underestimate heritability when intra-individual variability is moderate to high (which is common in many real-world scenarios). We supported this finding with analytical derivations, simulations and an experimental dataset from the ABCD study, validating the performance of the HLM approach. Our simulation results suggest that traits with small effect sizes may require much larger sample sizes than currently practiced.

## Acknowledgments

GC and PT were supported by the NIMH Intramural Research Program (ZICMH002888) of the NIH/HHS, USA. DM was also supported by the NIMH Intramural Research Program (ZICMH002960) of the NIH/HHS, USA. Data used in the preparation of this article were obtained from the Adolescent Brain Cognitive Development (ABCD) Study (https://abcdstudy.org), held in the NIMH Data Archive (NDA).

## Appendices

### A Simulation specifications

Simulations were conducted with the following specifications under the HLM formulation (20) compared to the conventional SEM (2). All distributions involved were assumed to be Gaussian, and the data were aggregated across trials for each condition when applied to the SEM formulation. While the Gaussian distribution is adopted here for simulation convenience and may not always be applicable in real-world scenarios, the simulation results still provide valuable insights into the impact of intra-individual variability and the necessary sample size requirements. The two overall effects under the HLM (20) were fixed at *α*_*c*_ = 0 (*c* = *c*_1_, *c*_2_). Since the standard deviation is essentially a scaling factor, the sum of inter-individual and inter-family variances was fixed as 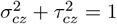 (*c* = *c*_1_, *c*_2_; *z* = MZ, DZ) without loss of generality. Four factors were changed to create the set of simulations across the following ranges of values:

(a) Three family sample sizes *F* = 100, 500, 1000, half of which were MZ (and DZ) zygosities.
(b) Three trial sample sizes *T* = 50, 100, 500.
(c) Three ratios of *R*_*v*_ = 1, 4, 10, leading to 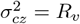.
(d) Two modeling approaches: SEM and HLM.

Each of these 3 *×* 3 *×* 3 *×* 2 = 54 combinations was simulated with 200 repetitions. For SEM, the function `lme` in the `R` package `nlme` (Pinheiro et al., 2022) was used to estimate variances. For HLM, the function `lmer` in the `R` package `lme4` (Bates et al., 2015) was adopted with the following iterative steps. The condition contrast was parameterized through the indicator variable defined in (22) in Subsection 3.5.

1) For each twin pair (*i*_1_, *i*_2_) within family *f* of zygosity *z*, obtain their effects at the two conditions *c*1 and *c*_2_ through randomly sampling from the following quadrivariate distribution:

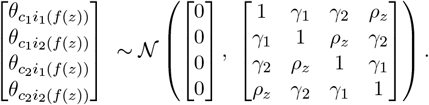 Here, *γ*_1_ and *γ*_2_ were fixed at 0.8 and 0.5, respectively, and *ρ*_*z*_ was chosen as 0.375 when *z* = MZ and 0.5 when *z* = DZ, so that the simulated correlation *rz* between a twin pair within a family regarding the condition contrast would be 0.85 for *z* = MZ and 0.6 for *z* = DZ, respectively. Thus, the targeted heritability *h*^2^ would be 0.5 per Falconer’s formula (6), while *c*^2^ and *e*^2^ would be 0.35 and 0.15.
2) Randomly draw trial-level data *y*_*ci*(*f* (*z*))_ from *N(α*_*c*_ + *θ*_*ci*(*f* (*z*))_, 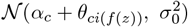) (*c* = *c*_1_, *c*_2_), where 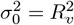.
3) Estimate 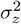 and 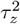. For SEM, obtain the average across trials for each condition and for each individual, and then apply the individual-level contrast between the conditions to the formulation (7) using the function `lme` from the `R` package `nlme`. For HLM, apply the model (20) using the function `lmer` from the `R` package `lme4`.
4) Obtain *r*_*z*_ using the formula (21). In addition, make two adjustments using formulas (17) and (18) with 1*/T* replaced by 2*/T*.
5) Estimate *h*^2^, *c*^2^, and *e*^2^ using Falconer’s formula (6).
6) Estimate bias adjustments for SEM: apply (18) with an extra factor of 2 to estimate SEM1, and (17) with an extra factor of 2 to estimate SEM2.

The results are shown in Figs. 7 and 8. The simulation scripts can be found online at https://github.com/afni/apaper_heritability.

**Figure 7.**
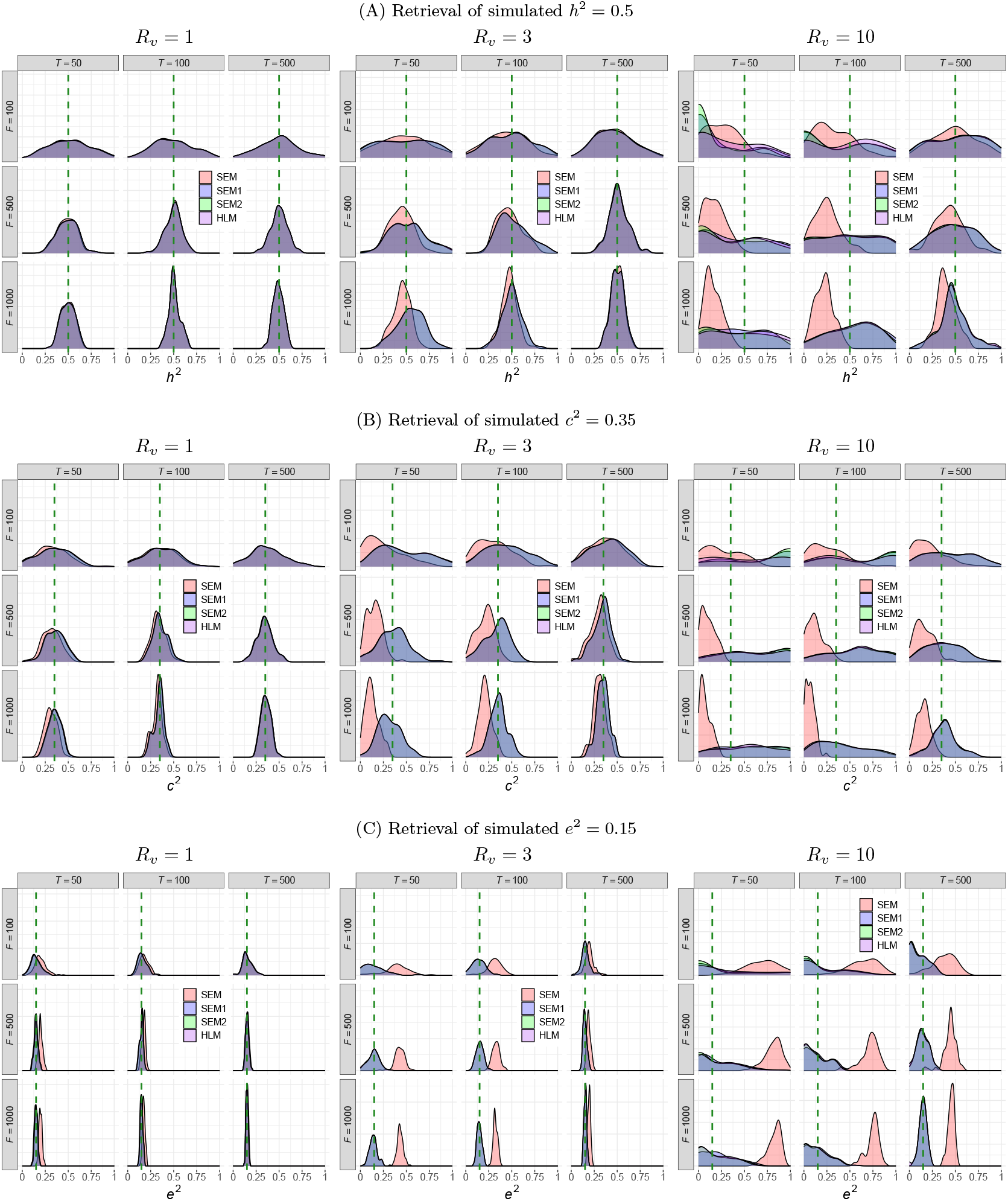
Simulation results of a trial-based experiment for the contrast between two conditions. Four factors were considered in the simulations: two models (SEM and HLM), three variability ratios (*R*_*v*_ = 1, 4, 10), three family sizes (*F* = 100, 500, 1000), and three trial sample sizes (*T* = 50, 100, 500). SEM1 and SEM2 represent the adjusted estimates obtained through the decontamination formulas (18) and (17), respectively. Each curve represents the density of parameter estimation from 200 repetitions, and each vertical dashed line indicates the simulated parameter value. The curves for HLM, SEM1, and SEM2 are nearly indistinguishable from each other.

**Figure 8.**
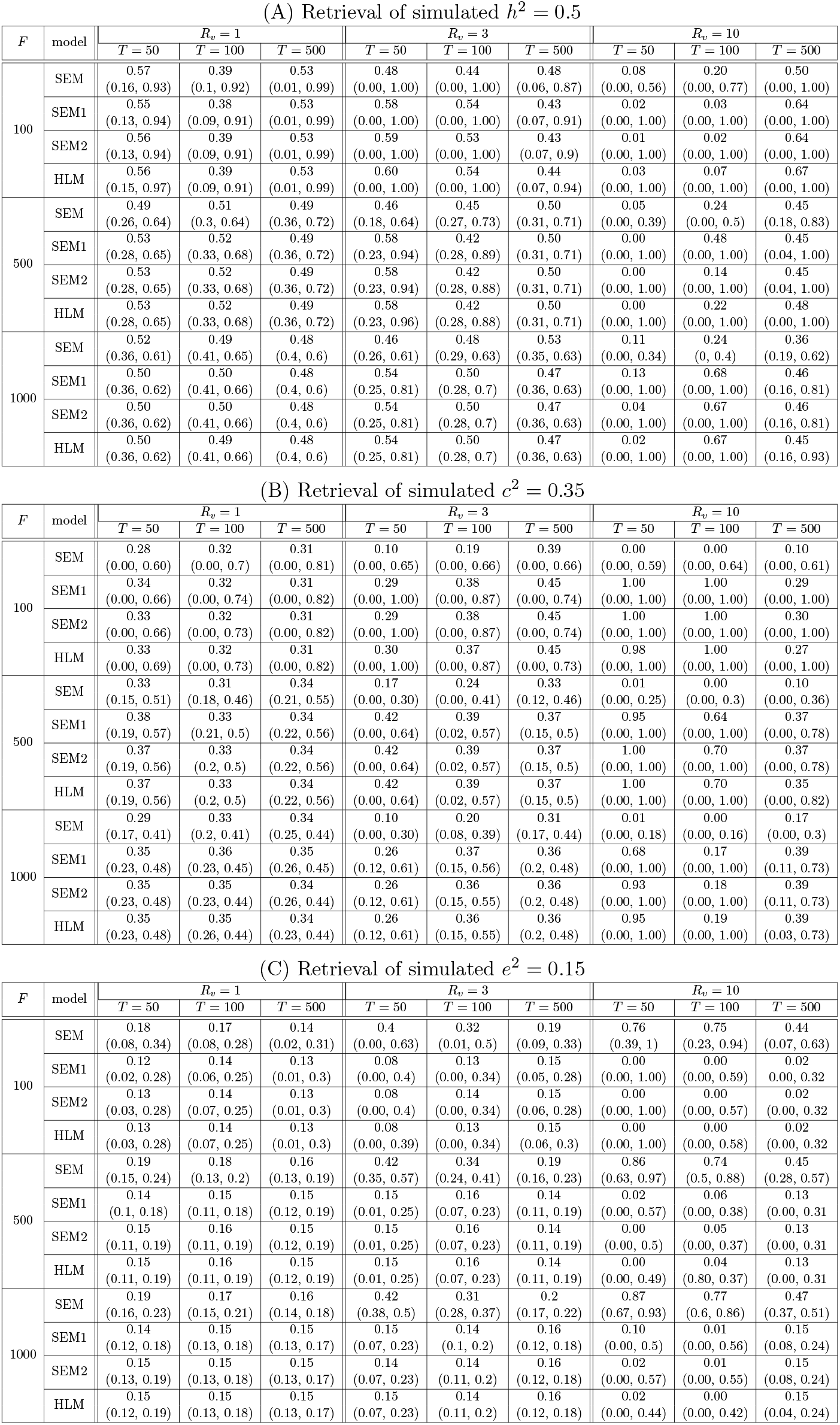
Summary of simulation results. The reported point estimates represent the modes, while the 95% uncertainty intervals, shown in parentheses, are highest density intervals derived from the simulations, as illustrated in Fig. 7.

## Footnotes

1 The term “uncertainty interval” is used here to include confidence intervals in frequentist usage and credible intervals in Bayesian methods.

